# Nonadditive gene expression is correlated with nonadditive phenotypic expression in interspecific triploid hybrids of willow (*Salix* spp.)

**DOI:** 10.1101/2021.10.13.464325

**Authors:** Craig H. Carlson, Yongwook Choi, Agnes P. Chan, Christopher D. Town, Lawrence B. Smart

## Abstract

Many studies have highlighted the complex and diverse basis for heterosis in inbred crops. Despite the lack of a consensus model, it is vital that we turn our attention to understanding heterosis in undomesticated, heterozygous, and polyploid species, such as willow (*Salix* spp.). Shrub willow is a dedicated energy crop bred to be fast-growing and high yielding on marginal land without competing with food crops. A trend in willow breeding is the consistent pattern of heterosis in triploids produced from crosses between diploid and tetraploid species. Here, we test whether differentially expressed genes are associated with heterosis in triploid families derived from diploid *S. purpurea*, diploid *S. viminalis*, and tetraploid *S. miyabeana* parents. Three biological replicates of shoot tips from all family progeny and parents were collected after 12 weeks in the greenhouse and RNA extracted for RNA-Seq analysis. This study provides evidence that nonadditive patterns of gene expression are correlated with nonadditive phenotypic expression in interspecific triploid hybrids of willow. Expression-level dominance was most correlated with heterosis for biomass yield traits and was highly enriched for processes involved in starch and sucrose metabolism. In addition, there was a global dosage effect of parent alleles in triploid hybrids, with expression proportional to copy number variation. Importantly, differentially expressed genes between family parents were most predictive of heterosis for both field and greenhouse collected traits. Altogether, these data will be used to progress models of heterosis to complement the growing genomic resources available for the improvement of heterozygous perennial bioenergy crops.

## INTRODUCTION

The heritability of gene expression has been attributed to both local *cis*-regulatory elements and distant *trans*-regulatory factors in the cell. Variation in these gene regulators can play dramatic roles in the evolution of gene expression. *Cis*-regulatory variation is thought to account for evolutionarily significant phenotypic differences, whereas *trans*-regulatory variation is thought to account more for adaptive differences (Wray 2007). For instance, *cis*-regulatory variation in promoter regions within-species should be minimal, compared to that among species. So, it is more likely that *trans*-effects should account for most of the regulatory variation in the intraspecific hybrid, and *cis*-effects in the interspecific hybrid (Wittkopp *et al*. 2008a). More simply, the greater phylogenetic distance between parents, the more likely it is that differential gene expression in the progeny will be due to gene localized polymorphism. However, it is uncertain whether nonadditive gene expression and regulatory divergence are more commonly observed in plants exhibiting hybrid vigor (heterosis).

Many studies on heterosis have focused on hybrids derived from crossing inbred parents (Guo *et al*. 2004; Guo *et al*. 2006), few on those derived from outcrossing parents (Landry *et al*. 2005; Zhuang and Adams 2007), and even fewer on hybrids derived from outcrossing parents of different species or ploidy (Wittkopp and Kalay 2011). From early expression studies based on only a few dozen genes to recent research employing RNA-Seq, a common result in maize, wheat, and rice is that additive gene expression in hybrids makes up the greatest proportion of those differentially expressed between the parents (Guo *et al*. 2006; Stupar and Springer 2006; Stupar *et al*. 2008; Wei *et al*. 2009), yet genes with nonadditive expression display allele-specific expression (ASE) (Guo *et al*. 2004; Springer and Stupar 2007; Wei *et al*. 2009). This differential expression could be due to the presence of remote *trans*-factors, whereby a small number of key regulatory genes could play significant roles in heterosis (Ni *et al*. 2009; He *et al*. 2010; Goff 2011).

A major ongoing topic in heterosis research is *to what extent is nonadditive gene expression correlated with nonadditive phenotypic expression* (Birchler *et al*. 2007), and how this informs combining ability or response to hybridization in the F_1_. In most crop plants, heterosis has been observed in hybrids bred from inbred parents of contrasting genetic backgrounds (East 1936; Birchler *et al*. 2003). In maize, high numbers of low-frequency alleles near conserved *cis*-regulatory regions in the genome have been thought to lead to gene misexpression and are implicated as having a deleterious impact on important component traits (Kremling *et al*. 2018). Dominance may help explain the phenomenon of heterosis in maize (McMullen *et al*. 2009), and there are efforts to purge these deleterious alleles from breeding material via targeted gene-editing technologies. Answers to questions regarding the genomic basis of heterosis are not only relevant in breeding and selection, but will contribute to our understanding of the evolution of dioecious plant species that regularly undergo interspecific hybridization and polyploidization events.

Willow (*Salix* spp.) is exceptionally diverse, with over 350 species characterized across most of the temperate range, and ploidy levels ranging from diploid to dodecaploid (Kuzovkina *et al*. 2008), so the genomic basis of heterosis is likely to be different from that of conventional crop plants. Interspecific hybridization has been a key component in shrub willow improvement, as F_1_ hybrids often display heterosis for biomass yield (Kopp *et al*. 2001; Cameron *et al*. 2008; Serapiglia *et al*. 2014a; Fabio *et al*. 2016), especially those derived from diploid and tetraploid parents (Smart and Cameron 2008; Serapiglia *et al*. 2014b; Carlson and Smart 2016). What is promising for the biomass production industry, is that these high-yielding triploids outperform foundational commercial cultivars for dry weight biomass yield and other biomass-related morphological and physiological traits (Fabio *et al*. 2017). While there is good evidence of heterosis in triploid hybrids of willow (Serapiglia *et al*. 2014a; Carlson and Smart 2021), the genomic basis of this phenomenon is not well-characterized.

To support breeding efforts, an Illumina-based reference genome assembly of female *S. purpurea* 94006 was constructed using a F_2_ map-guided approach to orient scaffolds into pseudomolecules (*Salix purpurea* v1.0, DOE-JGI, phytozome-next.jgi.doe.gov/). Recently, sex specific long-read genome assemblies of *S. purpurea* have been completed (*Salix purpurea* v5.1, *S. purpurea* ‘Fish Creek’ v.3.1, DOE-JGI, phytozome-next.jgi.doe.gov/). The genome size is an estimated 330 Mb and contains approximately 35,125 protein-coding genes (57,462 transcripts), and has proven useful in read alignment, variant discovery, and candidate gene selection (Hyden *et al*., 2021). There have been a handful of studies in shrub willow that focused on genetic mapping (Gunter *et al*. 2003; Berlin *et al*. 2010; Hanley and Karp 2016; Hällingback *et al*. 2016; Zhou *et al*. 2018; Carlson *et al*. 2019) of quantitative trait loci (QTL) associated with biomass yield traits to aid in marker-assisted selection (MAS), but most have been low-resolution. There have also been attempts at correlating cell wall biosynthesis genes with variation in biomass composition in *Salix* spp. (Serapiglia *et al*. 2012), as well as correlating sex dimorphism (Gouker *et al*. 2021) with gene expression and methylation patterns in F_2_ *S. purpurea* (Hyden *et al*. 2021). Thus far, family-based ASE in *Salix* is restricted to a single study of F_1_ and F_2_ intraspecific *S. purpurea* (Carlson *et al*. 2017), where expression-level dominance comprised the greatest proportion of differentially expressed genes between the parents of both families. Overall, there were more genes with ASE in the F_1_ compared to F_2_, but both families displayed greater levels of *cis*-than *trans*-regulatory divergent expression patterns. In high-yielding, triploid hybrids of bioenergy willow, the heritability of gene expression and its broad influence on modulating heterosis for biomass yield and other traits important for biomass production has not been characterized.

Using willow as a model for understanding heterosis in heterozygous polyploid perennials, the objectives of this study were to: 1) describe the inheritance and regulatory divergence patterns influencing gene expression within and among three interspecific hybrid triploid families, 2) test for dosage effects on parent alleles in triploid progeny, and 3) determine which genes and gene sets are most predictive of heterosis for biomass growth and wood chemical composition traits important for bioenergy production.

## MATERIALS AND METHODS

### Plant material and growing conditions

Progeny individuals from three full-sib F_1_ triploid families included in this study were derived from the interspecific crosses: *S. purpurea* 94006 × *S. miyabeana* 01-200-003 (Family 415), *S. viminalis* 07-MGB-5027 × *S. miyabeana* 01-200-003 (Family 423), and *S. miyabeana* 01-200-006 × *S. viminalis* ‘Jorr’ (Family 430). Herein, we refer to parents of the F_1_ families by their clone identifiers and discriminate the female and male parents as P1 and P2, respectively.

The field trial was established May 2014 at Cornell AgriTech (Geneva, NY). All parents and progeny were transplanted from nursery beds as stem cuttings (20 cm) in a randomized complete block design with four replicate blocks. The field perimeter was buffered using *S. purpurea* genotypes 94006 and ‘Fish Creek’ to avoid edge effects. Each plot consisted of three clones (within-row spacing: 0.4 m; between row spacing: 1.82 m), of which the middle plant was measured. The field trial was evaluated for three years.

Parent genotypes and randomly selected progeny were grown from stem cuttings (20 cm) in 12-L plastic pots with peat moss-based potting mix (Fafard, Agawam, MA) to evaluate growth traits under greenhouse conditions over the course of 12 weeks. Plot was defined as a single cutting planted in a pot, which were arranged in a randomized complete block design with four replicate blocks. Two blocks were located on benches in one greenhouse with the other two blocks in an adjacent greenhouse set for identical growing conditions. Supplemental greenhouse lighting was provided on a 14 hr day : 10 hr night regimen with maximum daytime temperature of 26° and a nighttime temperature of 18°. Beyond weekly applications of beneficial insects and mites for pest management, no pesticides were required, as there were no symptoms of biotic or abiotic stress on any plant material throughout the length of the study. Liquid fertilizer (Peter’s 15-16-17 Peat-Lite Special^®^, Scott’s, Marysville, OH) was applied weekly beginning four weeks after planting.

For more information on the experimental design and phenotypes recorded in the field and greenhouse trial, see Carlson and Smart (2021).

### Determination of ploidy level

The relative DNA content (pg 2C^-1^) of family parents and progeny was determined by flow cytometry using young leaf material harvested from actively growing shoots in greenhouse conditions. Analysis of 50 mg of mature leaf tissue from parental genotypes and selected progeny was performed at the Flow Cytometry and Imaging Core Laboratory at Virginia Mason Research Center in Seattle, WA. A minimum of four replicates of all samples were independently assessed using the diploid female *S. purpurea* clone 94006 as an internal standard. Diploid parent clones from multiple runs were averaged and then divided by the 2C-value of the check for that run. This factor was then multiplied by each sample value within the same run as the check. When a clone was analyzed more than once, 2C-values were averaged.

### Sample preparation and sequencing

A total of three biological replicate shoot tips (∼1 cm) of all triploid progeny individuals, as well as their parents, were excised from the primary stem and immediately flash-frozen in liquid N_2_ in the greenhouse, then placed in −80° storage. Shoot tips were defined as the shoot axis that is the most distal part of a shoot system, comprised of a shoot apical meristem and the youngest leaf primordia. For each sample, a single shoot tip was removed from −80° storage, and ground to a fine powder (100-200 mg) prior to RNA isolation using the Spectrum^™^ Total Plant RNA Kit with DNase I digestion (Sigma, St. Louis, MO). The only modification to manufacturer’s ‘Protocol B’ was that prior to the tissue lysis step, the 2-ME/lysate mixture was incubated at 65° for 5 min, otherwise, the manufacturers’ procedures were followed. After elution, cold ethanol precipitations were performed by the addition of 10 μL acetic acid and 280 μL 100% cold ethanol to 100 μL eluate and placed in −80° for 3 h. Samples were centrifuged at 17,000 × *g* for 30 min at 4°, washed with 80% ethanol, then centrifuged at 17,000 × *g* for 20 min at 4°. After centrifugation, the supernatant was discarded, and the pellet resuspended in ribonuclease-free 10 mM Tris-HCl. Quantification of RNA sample quality and concentration was performed using the Experion ‘StdSens’ kit (Bio-Rad Laboratories, Inc., Hercules, CA). Stranded RNA-Seq libraries were created and quantified by qPCR (2×76 bp or 2×151 bp) and sequenced on an Illumina Hi-Seq 2500 at J. Craig Venter Institute. Library sizes ranged from 8.3 to 53 million reads.

### Read filtering, mapping, and variant discovery

Low-coverage paired-end genomic DNA sequencing of the parents of the F_1_ families was performed to validate variants from RNA-Seq data. Biallelic SNPs were used to quantify allele-specific expression (ASE) within and among triploid progeny individuals. Parent DNA libraries were sequenced (Illumina HiSeq 2500, 2×101 bp) and aligned to the *S. purpurea* v1 reference genome using BWA mem (Li and Durbin 2009). Subsequent BAM files were sorted, marked for duplicates, and indexed in Picard (broadinstitute.github.io/picard). Indel realignment and variant calling was performed using *HaplotypeCaller* (emit_conf=10, call_conf=30) in the Genome Analysis Toolkit (GATK) (DePristo *et al*. 2011). Using *BBDuk* in the BBTools program (https://jgi.doe.gov/data-and-tools/bbtools/), raw reads were evaluated for artifact sequences by kmer matching (kmer = 25), allowing 1 mismatch and detected artifact was trimmed from the 3’ end of the reads. RNA spike-in reads, PhiX reads and reads containing any Ns were removed. Following quality trimming (phred = Q6), reads under the length threshold were removed (≥ 25 bp or 1/3 original read length). BWA mem was used for alignment of interleaved RNA-Seq reads to the reference. SAMtools was used to filter (-Shb -F 4 -f 0×2 -q 30), sort, and index resulting sequence alignment files. Duplicate reads were flagged using *MarkDuplicates* in Picard and GATK was used to flag and realign indels with *RealignmentTargetCreator* (minReads = 20) and *IndelRealigner*.

### Gene expression inheritance classifications

To categorize inheritance of gene expression in the hybrid, Negative Binomial (NB) exact tests were performed in edgeR (Robinson *et al*. 2010) in R (R Core Team 2021), at a False Discovery Rate (FDR) of 0.005, for only genes with a minimum counts-per-million (CPM) ≥ 1. Prior to NB tests, dispersions were estimated using three biological replicates of each group to account for library-to-library variability. Tests for differential expression were for paired comparisons between 1) diploid and tetraploid parents (P_2X_ and P_4X_), 2) diploid parent and the triploid hybrid (P_2X_ and H), and 3) tetraploid parent and the triploid hybrid (P_4X_ and H). Gene expression inheritance classifications were based on log_2_ fold-change > 1.2 and *q*-value < 0.005 resulting from exact tests between the parents and hybrid, according to Carlson et al. (2017).

### Regulatory divergence classifications

To determine *cis*- and *trans*-effects on gene expression, separate binomial exact tests were performed using library-normalized read counts of diploid (P_2X_) and tetraploid (P_4X_) parent alleles in the parents (P_2X_ and P_4X_) and the F_1_ triploid progeny individual (H_2X_ and H_4X_) from the P_2X_ × P_4X_ cross. For a two-sided binomial test, the null hypothesis is that the expected counts are in the same proportions as the library sizes, or that the binomial probability for the first library is *n*_1_ / (*n*_1_ + *n*_2_). To test the null of independence of rows (P_2X_ vs P_4X_ and H_2X_ vs H_4X_) and columns (P_2X_ vs H_2X_ and P_4X_ vs H_4X_), Fisher’s exact test was performed on a 2×2 matrix comprised of P_2X_ and P_4X_ and H_2X_ and H_4X_ normalized read counts. For all tests, a fixed FDR was applied at a level of 0.005. Filtering parameters required ≥ 20 reads summed between the parents, and two alleles at a locus, such that each allele corresponds to either the diploid or tetraploid parent.

For each site, significant differences (FDR = 0.005) on the expression of parent alleles can occur either between the parents (P, binomial exact test), the hybrid (H, binomial exact test), or all (F, Fisher’s exact test). Categories of regulatory functions considered *cis*-only, *trans*-only, *cis* + *trans*, *cis* × *trans*, compensatory were assigned following previously described methods (Landry *et al*. 2005; McManus *et al*. 2010). Conservation of expression was attributed to cases where no significant differences could be observed. Ambiguous cases were observed when only one of the three tests (P, H, or F, described above) were deemed significant. While ambiguous cases could somewhat be resolved by lowering the significance threshold (e.g., FDR = 0.05), approximately equal proportions of ambiguous assignments were observed across regulatory divergence classes and triploid individuals. However, parent-only (P)-ambiguous genes were more common than the other ambiguous cases, of which, F-ambiguous genes were the least frequent.

### Copy number variation

Copy number variation (CNV) was analyzed on a chromosome-wide scale, using median log_2_ (P_2X_ / P_4X_) difference of logs in the parents and the median percentage of reads attributable to the P_2X_ allele in the triploid hybrid (diploid %). Diploid % was calculated as H_P2X_ / (H_P2X_ + H_P4X_) × 100, where H_P2X_ is a vector of library-normalized counts of the P_2X_ allele in the hybrid and H_P4X_ is that of the P_4X_ allele in the hybrid. The expected CN of each homeolog in the hybrid was either determined to be deficient, normal, or replete, depending on these two parameters (Figure S1). To avoid over-estimating CNV in triploids, binned coverage of paired-end Illumina DNA-Seq reads of the parents was compared to validate RNA-Seq results.

### Gene ontology analysis

Gene ontology (GO) term enrichment was performed in agriGO (Du *et al*. 2010) using the subset of the *S. purpurea* v1 transcriptome (reference set) that passed filtering, prior to tests of differential expression. Only significant ontologies (FDR = 0.05) were reported. *Salix purpurea* gene models and associated GO-terms which were annotated as hypothetical proteins were inferred using the best-hit (blastp e-value ≤ 0.01) to *Populus trichocarpa* (Phytozome v10.3) and *Arabidopsis thaliana* (TAIR10 and Araport11 annotations) proteomes.

### Gene-trait correlations and prediction of heterosis using selected and random gene sets

Gene-trait correlations were performed for each family using scaled log_2_ (CPM + 1) library-normalized gene expression values (File S1) and midparent heterosis values for field and greenhouse collected traits (File S2), which were calculated as the percent deviation of the F_1_ from the midparent value, as described in Carlson and Smart (2021). Ridge regression (α = 0) was used to predict midparent heterosis trait values with different gene sets using 10 replications of nested cross validation (tenfold inner and outer) with *cv.glmnet* in glmnet (Friedman *et al*. 2010). Gene sets were comprised of scaled log_2_ (CPM + 1) library normalized expression values of: 1) 5,000 randomly sampled genes, 2) the top 5,000 most highly expressed genes, 3) 4,986 genes which were differentially expressed between at least one pair of family parents, and 4) 379 genes that were commonly differentially expressed between all three family parent pairs. Prediction accuracy was assessed via linear regression of mean predicted and observed values.

## RESULTS

### Transcriptome analysis

After quality filtering and alignment of triploid F_1_ progeny and parent paired-end RNA-Seq reads to the *S. purpurea* v1 reference, the library sizes ranged from 10 to 56 million. Of the 105 libraries sequenced, there were two identified as outliers (13X-430-035, greenhouse plot 59, biological replicate 1; 12X-415-074, greenhouse plot 278, biological replicate 3) and removed prior to downstream statistical analyses. In a multi-dimensional scaling plot of normalized transcriptome-wide gene expression of all triploid F_1_ progeny individuals and their parents (Figure 1A), the first dimension represents sample distances based on species pedigree (Ve = 27%), with individuals containing *S. viminalis* in their background clustering to the right of the first dimension, *S. miyabeana* in the center, and *S. purpurea* to the left, such that the *S. viminalis* parents (07-MBG-5027 and ‘Jorr’) and *S. purpurea* 94006 are at extremes, or most distantly related. While family 415 and 423 individuals share the common tetraploid *S. miyabeana* parent 01-200-003, the proximity of family 423 and 430 clusters indicates that common parent species (*S. viminalis* and *S. miyabeana*) is a more important factor contributing to transcriptome-wide distances. The second and third dimensions further separate sample libraries by pedigree (Ve = 11%) and ploidy (Ve = 7%), respectively (Figure 1C). For all three triploid families, the respective diploid and tetraploid parents flank clusters of family individuals and are relatively equidistant from the offspring cluster centers. Taking the first two dimensions into account, Euclidean distances approximated here implies transcriptome-wide gene expression inheritance is mostly conserved or additive.

**Figure 1.**
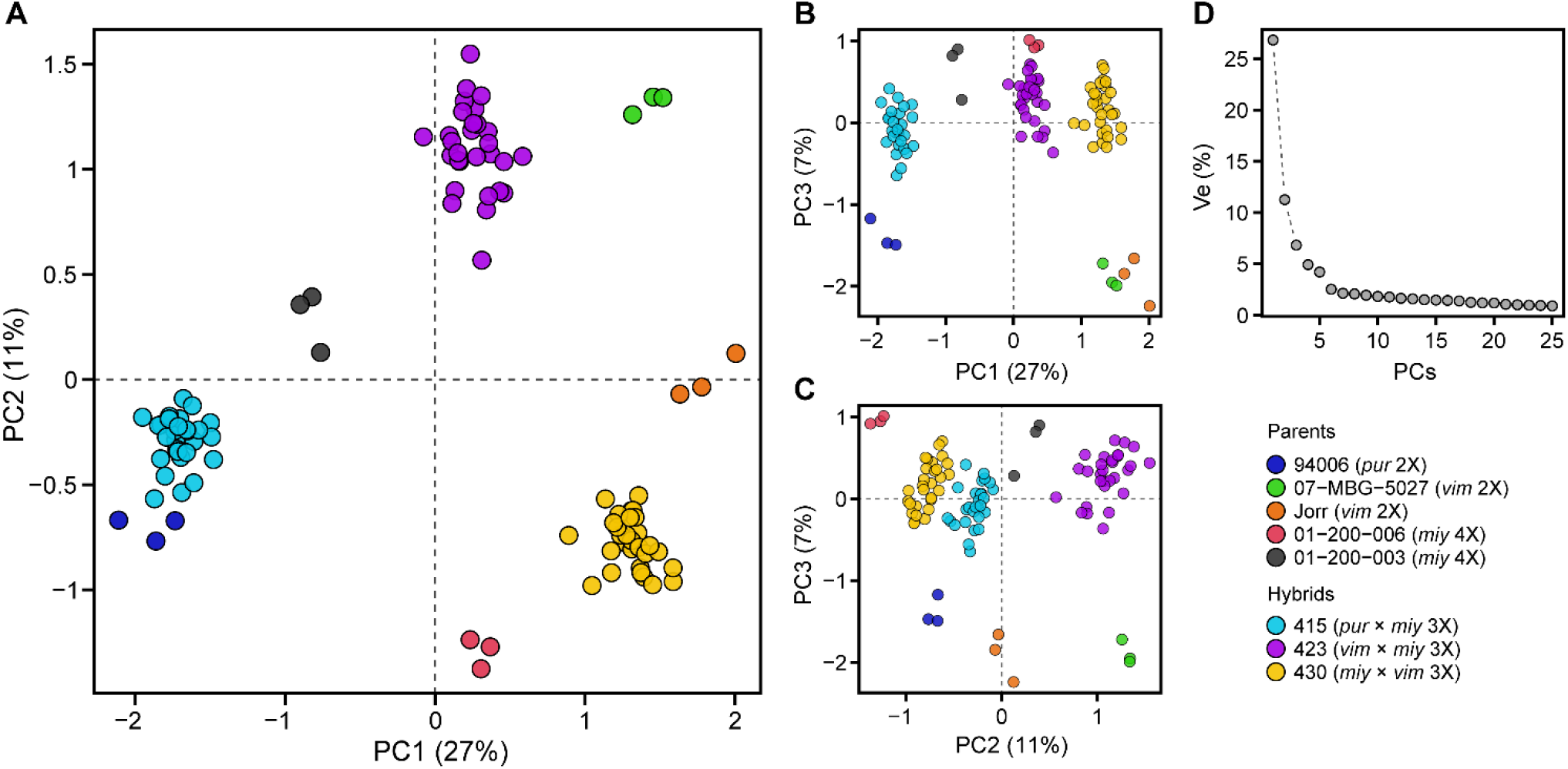
Multi-dimensional scaling plot of library-normalized transcriptome-wide gene expression of all triploid F_1_ progeny individuals (families 415, 423, and 430) and their diploid (94006, 07-MBG-5027, and ‘Jorr’) and tetraploid (01-200-006 and 01-200-003) parents. Panel **(A)** PC1 versus PC2, **(B)** PC1 versus PC3, **(C)** PC2 versus PC3, and **(D)** percent variance explained (% Ve) by the first 25 PCs. Euclidean distances on the two-dimensional plot approximate leading log_2_ fold-changes between samples, using the top 500 genes with the largest standard deviations. Parents and progeny libraries are colored according to the legend.

### Differential gene expression

Exact tests (FDR = 0.005) of differential gene expression between triploid family parent genotypes yielded similar numbers of differentially expressed genes, but the P1:P2 ratios differed (Table 1). The comparison of family 415 parents, *S. purpurea* 94006 (P1) versus *S. miyabeana* 01-200-003 (P2), had 5,166 differentially expressed genes, with 2,661 genes greater in 94006 and 2,505 genes greater in 01-200-003 (P1:P2 = 1.06). The family 423 parents, *S. viminalis* 07-MBG-5027 (P1) versus *S. miyabeana* 01-200-003 (P2), had 5,523 differentially expressed genes, with 2,469 genes greater in 07-MBG-5027 and 3,054 genes higher expressed in 01-200-003 (P1:P2 = 0.81). The family 430 parent comparison, *S. miyabeana* 01-200-006 (P1) versus *S. viminalis* ‘Jorr’ (P2), yielded 5,155 differentially expressed genes, with 2,467 genes greater in 01-200-006 and 2,688 genes greater in Jorr (P1:P2 = 0.91). Globally, the parents of family 423 had a greater percentage of genes that were differentially expressed (22.1%), compared to the parents of families 415 (20.8%) and 430 (20.5%). A total of 379 genes were differentially expressed in common among all three family parent duos.

**Table 1.** Number of differentially expressed genes between triploid family parents.

| Family | Female (P1) parent | Male (P2) parent | P1 > P2 (%) | P1 < P2 (%) | P1 = P2 (%) | Total |
| --- | --- | --- | --- | --- | --- | --- |
| 415 | 94006 [2x] | 01-200-003 [4x] | 2,661 (10.7) | 2,505 (10.1) | 19,641 (79.2) | 24,807 |
| 423 | 07-MBG-5027 [2x] | 01-200-003 [4x] | 2,469 (9.86) | 3,054 (12.2) | 19,519 (77.9) | 25,042 |
| 430 | 01-200-006 [4x] | Jorr [2x] | 2,467 (9.81) | 2,688 (10.7) | 19,993 (79.5) | 25,148 |

For those genes differentially expressed between parents, inheritance patterns were determined based on both the parent expression values and those observed in the hybrids (Table 2). The percentage of differentially expressed genes showing nonadditive inheritance ranged from 27% to 39% (mean = 33.5%) (Figure S2) in family 415, 40% to 56% (mean = 49.8%) in family 423, and 34% to 60% (mean = 50.3%) in family 430. Transgressively expressed genes (under- and overdominant) averaged just 0.7%, 1.1%, and 1.0%, for families 415, 423, and 430, respectively. The percentage of genes with underdominant expression out of total transgressively expressed genes was 98%, 74%, and 80% for families 423, 430, and 415, respectively. All individuals had a greater percentage of genes with expression level dominance in the direction of the tetraploid parent, ranging from 66% to 88% and a mean of 70% across all triploid families.

**Table 2.**
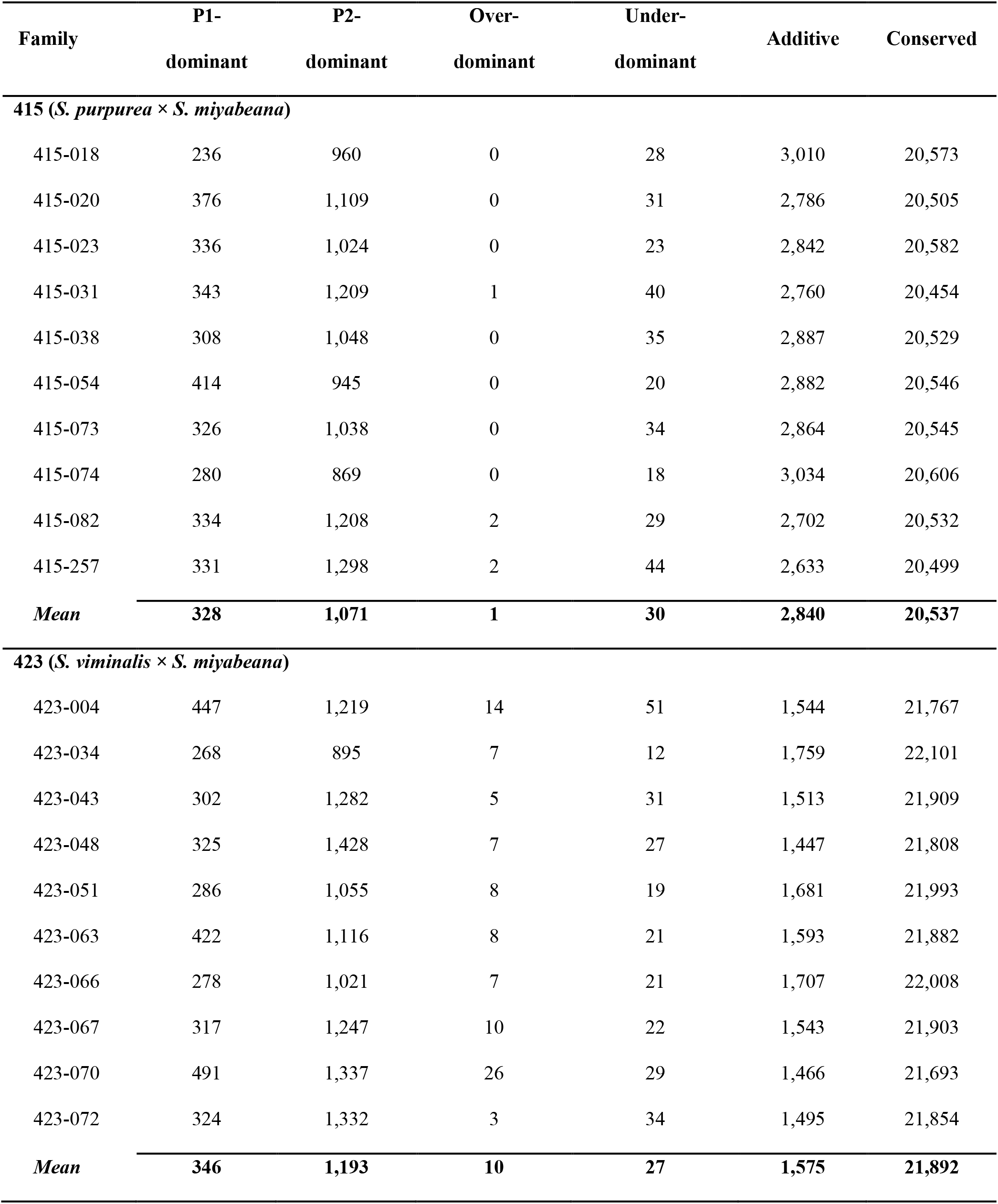
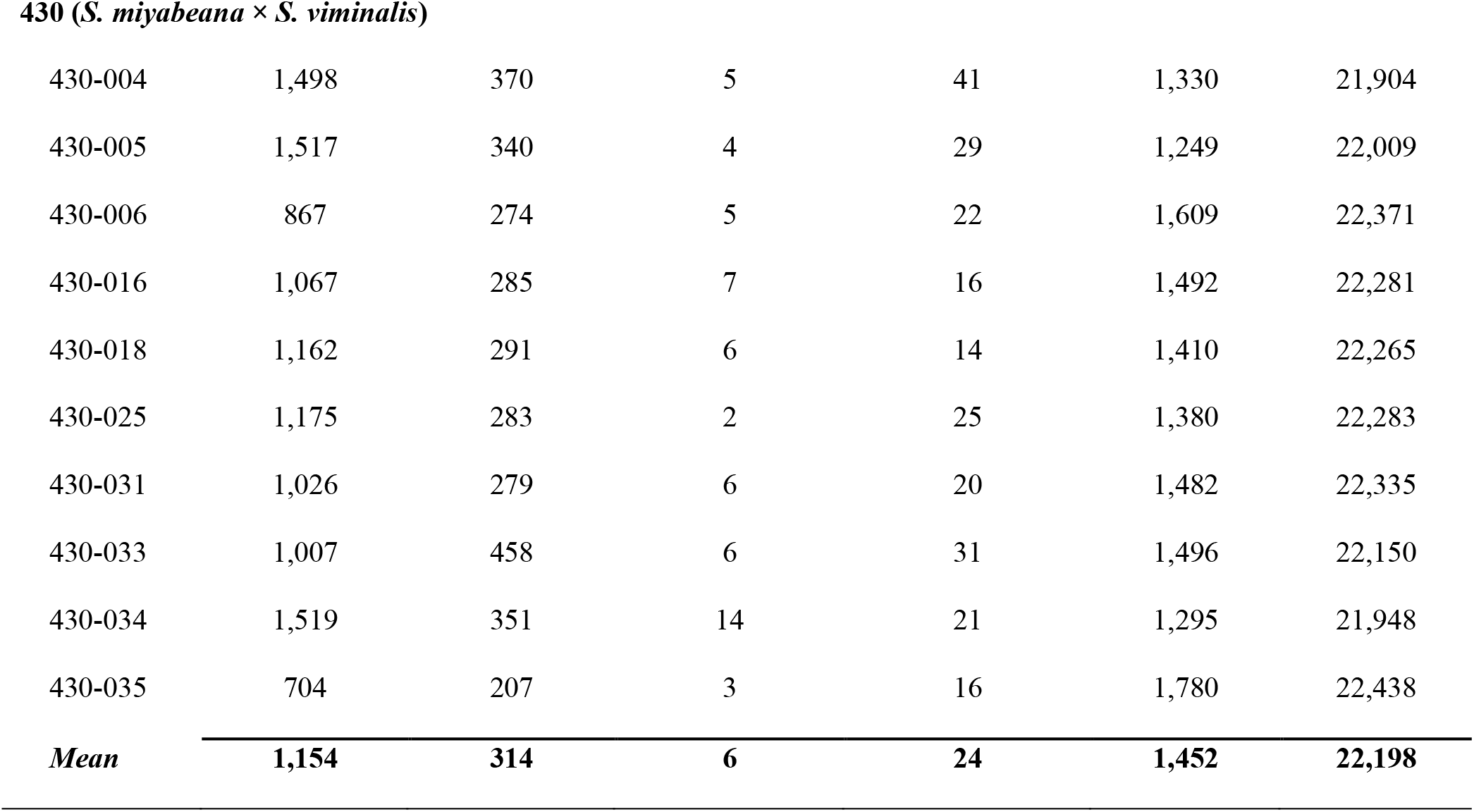
Number of genes assigned to inheritance classifications in triploid F_1_ progeny individuals and their averages by family.

| Family | P1-dominant | P2-dominant | Over-<br>dominant | Under-<br>dominant | Additive | Conserved |
| --- | --- | --- | --- | --- | --- | --- |
| **415 (*S. purpurea* × *S. miyabeana*)** |  |  |  |  |  |  |
| 415-018 | 236 | 960 | 0 | 28 | 3,010 | 20,573 |
| 415-020 | 376 | 1,109 | 0 | 31 | 2,786 | 20,505 |
| 415-023 | 336 | 1,024 | 0 | 23 | 2,842 | 20,582 |
| 415-031 | 343 | 1,209 | 1 | 40 | 2,760 | 20,454 |
| 415-038 | 308 | 1,048 | 0 | 35 | 2,887 | 20,529 |
| 415-054 | 414 | 945 | 0 | 20 | 2,882 | 20,546 |
| 415-073 | 326 | 1,038 | 0 | 34 | 2,864 | 20,545 |
| 415-074 | 280 | 869 | 0 | 18 | 3,034 | 20,606 |
| 415-082 | 334 | 1,208 | 2 | 29 | 2,702 | 20,532 |
| 415-257 | 331 | 1,298 | 2 | 44 | 2,633 | 20,499 |
| **Mean** | **328** | **1,071** | **1** | **30** | **2,840** | **20,537** |
| **423 (*S. viminalis* × *S. miyabeana*)** |  |  |  |  |  |  |
| 423-004 | 447 | 1,219 | 14 | 51 | 1,544 | 21,767 |
| 423-034 | 268 | 895 | 7 | 12 | 1,759 | 22,101 |
| 423-043 | 302 | 1,282 | 5 | 31 | 1,513 | 21,909 |
| 423-048 | 325 | 1,428 | 7 | 27 | 1,447 | 21,808 |
| 423-051 | 286 | 1,055 | 8 | 19 | 1,681 | 21,993 |
| 423-063 | 422 | 1,116 | 8 | 21 | 1,593 | 21,882 |
| 423-066 | 278 | 1,021 | 7 | 21 | 1,707 | 22,008 |
| 423-067 | 317 | 1,247 | 10 | 22 | 1,543 | 21,903 |
| 423-070 | 491 | 1,337 | 26 | 29 | 1,466 | 21,693 |
| 423-072 | 324 | 1,332 | 3 | 34 | 1,495 | 21,854 |
| **Mean** | **346** | **1,193** | **10** | **27** | **1,575** | **21,892** |
| **430 (*S. miyabeana* × *S. viminalis*)** |  |  |  |  |  |  |
| 430-004 | 1,498 | 370 | 5 | 41 | 1,330 | 21,904 |
| 430-005 | 1,517 | 340 | 4 | 29 | 1,249 | 22,009 |
| 430-006 | 867 | 274 | 5 | 22 | 1,609 | 22,371 |
| 430-016 | 1,067 | 285 | 7 | 16 | 1,492 | 22,281 |
| 430-018 | 1,162 | 291 | 6 | 14 | 1,410 | 22,265 |
| 430-025 | 1,175 | 283 | 2 | 25 | 1,380 | 22,283 |
| 430-031 | 1,026 | 279 | 6 | 20 | 1,482 | 22,335 |
| 430-033 | 1,007 | 458 | 6 | 31 | 1,496 | 22,150 |
| 430-034 | 1,519 | 351 | 14 | 21 | 1,295 | 21,948 |
| 430-035 | 704 | 207 | 3 | 16 | 1,780 | 22,438 |
| **Mean** | **1,154** | **314** | **6** | **24** | **1,452** | **22,198** |

There were fewer numbers of diploid parent dominant genes (15) (Table S1) than tetraploid parent dominant genes (89) (Table S2) that were common across all families and individuals. Due to the low number of common diploid parent dominant genes, there were no significant functional enrichments. Tetraploid dominant genes were enriched for GO molecular functions: beta-glucosidase activity and catalytic activity. In addition, tetraploid parent dominant genes were enriched for the KEGG pathways: phenylpropanoid biosynthesis, cyanoamino acid metabolism, biosynthesis of secondary metabolites, metabolic pathways, and starch and sucrose metabolism (Table S3).

### Allele-specific expression

To determine the extent of regulatory divergent expression, tests for ASE were conducted using expression data on biallelic sites that were first called with parent DNA-Seq and RNA-Seq libraries prior to calling parent alleles in the progeny. Family averages for the total number of genes assigned to at least one regulatory class were 15,391 (±114), 16,800 (±72), and 16,711 (±113), for families 415, 423, and 430, respectively (Table 3). On average, the percentage of genes assigned to non-conserved regulatory classes was 12%, 11%, and 10%, for families 415, 423, ad 430, respectively (Figure S3). Family 415 had the greatest percentage of non-conserved genes with *cis*-regulation (65%), compared to families 423 (58%) and 430 (54%). The greatest mean percentage of genes with *trans* (24.6%), *cis* × *trans* (7.4%), and compensatory (10.8%) regulatory divergence patterns was for family 430, whereas family 415 had the greatest mean *cis* + *trans* (5.1%). Across all triploid individuals, a total of 49 genes were in common, having either *cis*, *trans*, *cis* + *trans*, *cis* × *trans*, or compensatory regulatory classifications (Table S4). In addition, higher proportions of overdominant and underdominant expression coincided with higher proportions of *cis* × *trans* and compensatory regulatory classes. Further, a higher proportion of *cis* + *trans* divergence coincided with a lower proportion of underdominant expression, most notably for the comparison of families 423 and 430 with family 415.

**Table 3.**
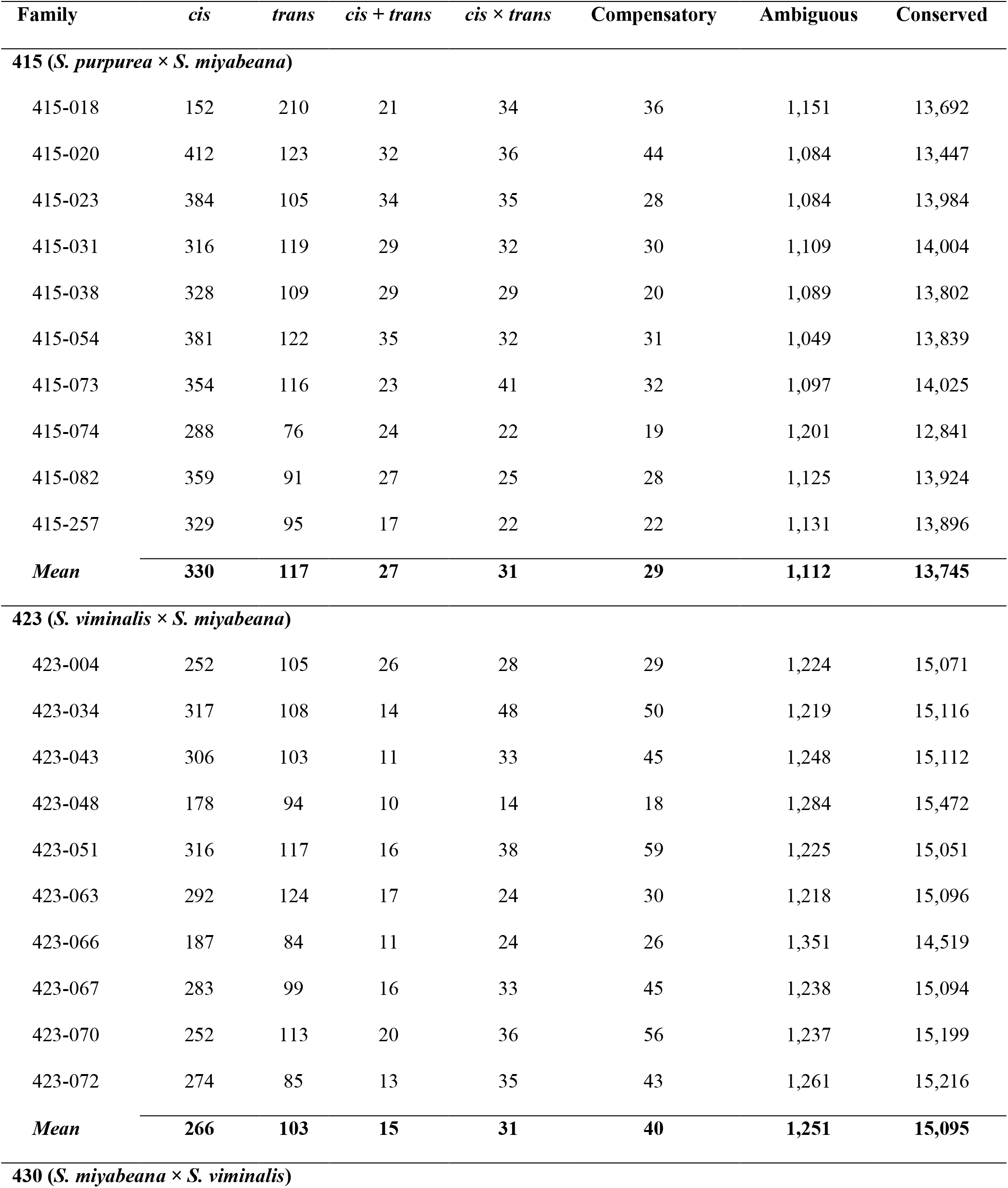
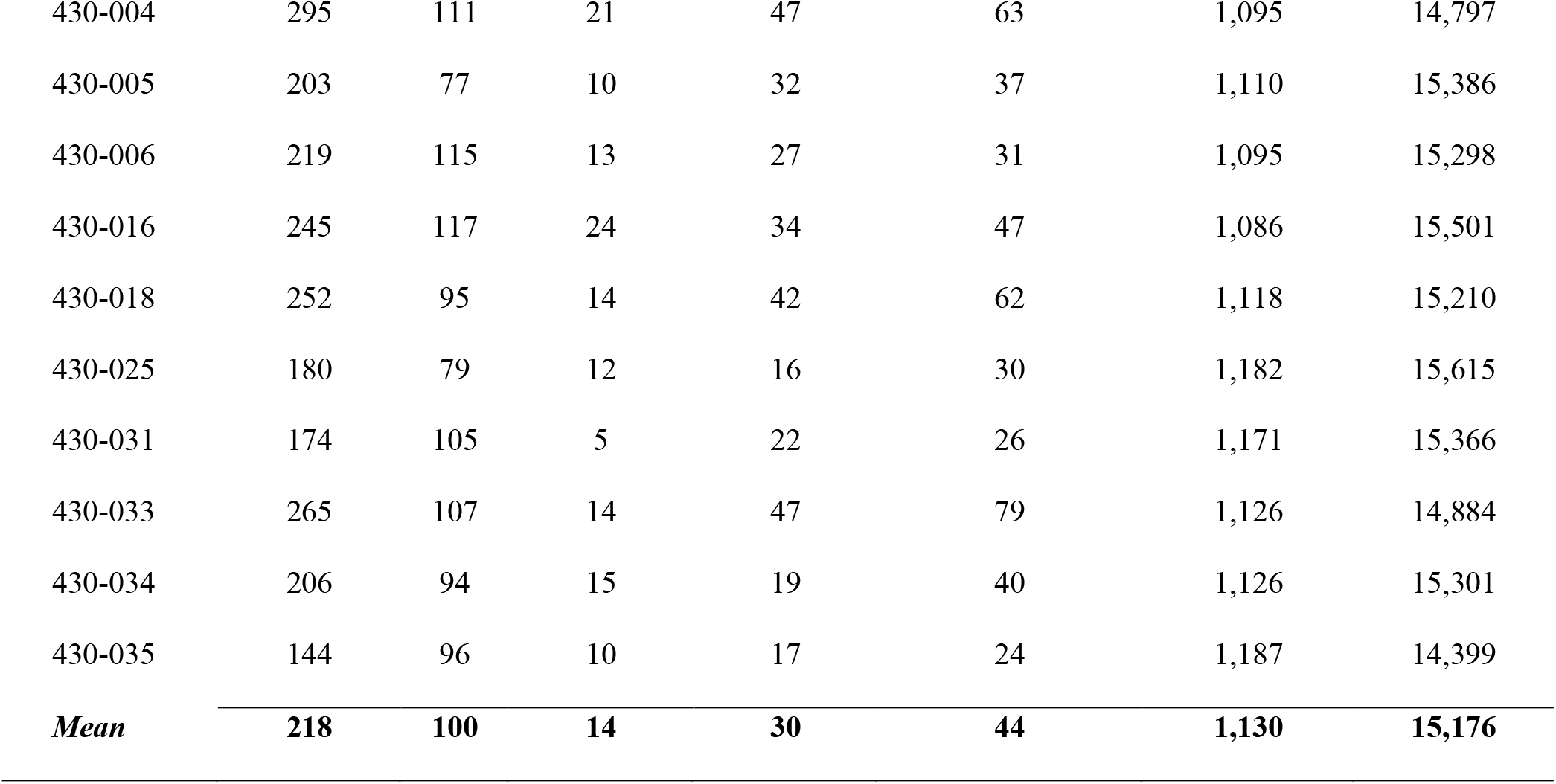
Number of genes assigned to regulatory divergence classifications (FDR = 0.005) in triploid F_1_ individuals and their means by family.

| Family | *cis* | *trans* | *cis* + *trans* | *cis* × *trans* | Compensatory | Ambiguous | Conserved |
| --- | --- | --- | --- | --- | --- | --- | --- |
| **415 (*S. purpurea* × *S. miyabeana*)** |  |  |  |  |  |  |  |
| 415-018 | 152 | 210 | 21 | 34 | 36 | 1,151 | 13,692 |
| 415-020 | 412 | 123 | 32 | 36 | 44 | 1,084 | 13,447 |
| 415-023 | 384 | 105 | 34 | 35 | 28 | 1,084 | 13,984 |
| 415-031 | 316 | 119 | 29 | 32 | 30 | 1,109 | 14,004 |
| 415-038 | 328 | 109 | 29 | 29 | 20 | 1,089 | 13,802 |
| 415-054 | 381 | 122 | 35 | 32 | 31 | 1,049 | 13,839 |
| 415-073 | 354 | 116 | 23 | 41 | 32 | 1,097 | 14,025 |
| 415-074 | 288 | 76 | 24 | 22 | 19 | 1,201 | 12,841 |
| 415-082 | 359 | 91 | 27 | 25 | 28 | 1,125 | 13,924 |
| 415-257 | 329 | 95 | 17 | 22 | 22 | 1,131 | 13,896 |
| **Mean** | **330** | **117** | **27** | **31** | **29** | **1,112** | **13,745** |
| **423 (*S. viminalis* × *S. miyabeana*)** |  |  |  |  |  |  |  |
| 423-004 | 252 | 105 | 26 | 28 | 29 | 1,224 | 15,071 |
| 423-034 | 317 | 108 | 14 | 48 | 50 | 1,219 | 15,116 |
| 423-043 | 306 | 103 | 11 | 33 | 45 | 1,248 | 15,112 |
| 423-048 | 178 | 94 | 10 | 14 | 18 | 1,284 | 15,472 |
| 423-051 | 316 | 117 | 16 | 38 | 59 | 1,225 | 15,051 |
| 423-063 | 292 | 124 | 17 | 24 | 30 | 1,218 | 15,096 |
| 423-066 | 187 | 84 | 11 | 24 | 26 | 1,351 | 14,519 |
| 423-067 | 283 | 99 | 16 | 33 | 45 | 1,238 | 15,094 |
| 423-070 | 252 | 113 | 20 | 36 | 56 | 1,237 | 15,199 |
| 423-072 | 274 | 85 | 13 | 35 | 43 | 1,261 | 15,216 |
| **Mean** | **266** | **103** | **15** | **31** | **40** | **1,251** | **15,095** |
| **430 (*S. miyabeana* × *S. viminalis*)** |  |  |  |  |  |  |  |
| 430-004 | 295 | 111 | 21 | 47 | 63 | 1,095 | 14,797 |
| 430-005 | 203 | 77 | 10 | 32 | 37 | 1,110 | 15,386 |
| 430-006 | 219 | 115 | 13 | 27 | 31 | 1,095 | 15,298 |
| 430-016 | 245 | 117 | 24 | 34 | 47 | 1,086 | 15,501 |
| 430-018 | 252 | 95 | 14 | 42 | 62 | 1,118 | 15,210 |
| 430-025 | 180 | 79 | 12 | 16 | 30 | 1,182 | 15,615 |
| 430-031 | 174 | 105 | 5 | 22 | 26 | 1,171 | 15,366 |
| 430-033 | 265 | 107 | 14 | 47 | 79 | 1,126 | 14,884 |
| 430-034 | 206 | 94 | 15 | 19 | 40 | 1,126 | 15,301 |
| 430-035 | 144 | 96 | 10 | 17 | 24 | 1,187 | 14,399 |
| *Mean* | **218** | **100** | **14** | **30** | **44** | **1,130** | **15,176** |

### Gene activation and silencing

The presence (CPM > 1) or absence (CPM = 0) of transcripts in the parent and triploid hybrid was compared for each family. Overall, more genes were silenced than activated in the triploid hybrids, especially for families 415 and 430, in which nearly five-times the number of genes were silenced than activated (Figure S4). Family 423 had a greater number of genes activated than the other two families, whereas family 430 had the greatest number of genes silenced. There were no GO-terms enriched for either activated or silenced gene-sets.

### Dosage effects on gene expression

To test whether there was a dosage effect on parent alleles in triploid progeny, ASE ratios were compared within and among families. Only extreme deviations from expected dosage ratios (Pr = 1×10^-5^) were included in the analysis and considered to be dysregulated. Since it is expected that the triploid hybrid has inherited a single copy of the diploid parent allele and two copies of the tetraploid parent allele, if there was no deviation in expression of the parent alleles in the hybrid, all loci would be represented by a single point at the intersection of expected P_2X_ / P_4X_ difference of logs, log_2_ (P_2X_ / P_4X_) (Figure 2).

**Figure 2.**
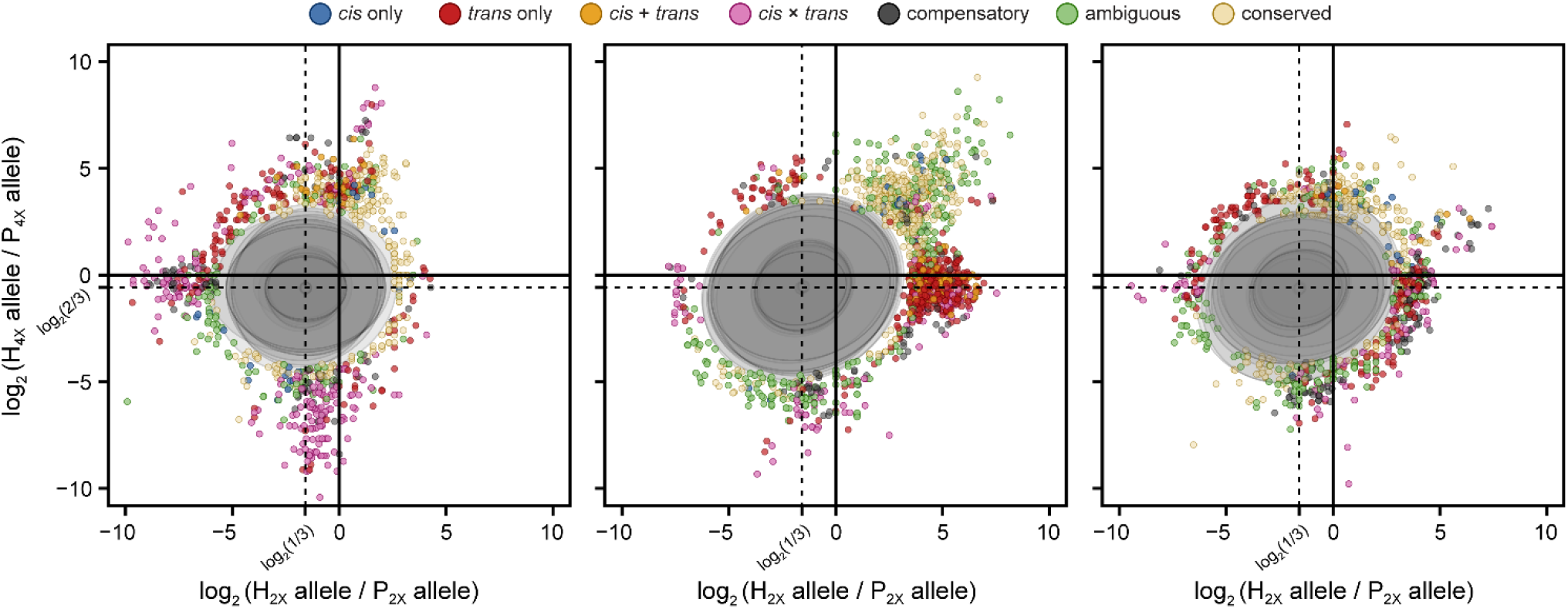
Superimposed dosage differential scatterplots of 10 individuals from each of the families 415, 423, and 430 (left to right, respectively). Each point depicts the log_2_ ratio of the diploid parent allele in the hybrid and the diploid parent allele in the parent against the log_2_ ratio of the tetraploid parent allele in the hybrid and the tetraploid parent allele in the parent. Points are colored according to their regulatory assignment. Ellipses mask most of the distribution of log_2_ dosage ratios (Pr = 1×10^-5^), such that points sitting outside ellipses are extreme outliers from expected dosage. Dotted lines at log_2_ (1/3) = −1.585 and log_2_ (2/3) = −0.585 represent distribution averages for diploid and tetraploid ratios, which is where the average distribution of dosage ratios is expected to occur.

While the dosage ratios were comparable within in families, and all genome-wide family means fell within expected ranges, there were significant departures from expected dosage. For dosage ratio outliers, all three triploid families exhibited unique patterns. There was an abundance of genes showing up- and down-regulation of the tetraploid parent allele in family 415, a majority of which showed *cis* × *trans* regulatory divergence. Family 423 outliers featured up-regulation of the diploid parent allele in the hybrid, as well as high expression levels of both diploid and tetraploid parent alleles in the hybrid (i.e., *trans*). Many outliers with greater diploid and tetraploid ASE in the hybrid were classified ambiguous, and those ratios were not different for the tetraploid parent allele, but a greater diploid ratio had either *trans* or *cis* × *trans* regulatory patterns. All regulatory patterns were represented in family 430, which had the greatest number of unique genes among the families that had dosage ratio outliers from at least one individual. Unlike family 423, there were few genes with greater expression for both diploid and tetraploid parents in family 430. In general, common dosage dysregulated genes showed significant enrichment for response to stress, transcription, small molecule activity, and binding activity (Table S5).

Genes with the greatest mean expression coincided with greater gene expression variance. Those genes with a μ/σ < 1 (n = 338) were significantly (*q*-value < 0.005) enriched for: GO biological functions of photosynthesis, translation, and response to stimulus; GO molecular functions of structural molecule activity, structural constituent of the ribosome, tetrapyrrole binding, and chlorophyll binding; GO cellular components of chloroplast, cytoplasm, chloroplast thylakoid, photosystem, and apoplast; KEGG pathways of photosynthesis, photosynthesis - antenna proteins, ribosome, metabolic pathways, and flavonoid biosynthesis. Over 50% of the top 50 most variable genes are either class I chaperonin heat shock proteins or ribosomal complex subunits, with the latter being most prominent. On the converse, the most highly expressed but *least* variable genes were enriched for the GO molecular function of ADP binding, of which most are annotated as stress-associated or disease resistance proteins (e.g., receptor-like kinases) and pentatricopeptide repeat-containing proteins.

### Chromosomal copy number variation

The difference in log_2_ (P_2X_ / P_4X_) expression of parent alleles in the hybrid from respective diploid and tetraploid progenitors can help determine if major departures from expected dosage in the hybrid are a result of copy number variation (CNV) in the tetraploid parent. For instance, it is expected that triploids inherit one chromosome copy from the diploid parent, and two copies from the tetraploid parent, such that the difference of the hybrid log_2_ (P_2X_ / P_4X_) from the parent is equal to 1. Although these ratios are tetraploid parent informative, aneuploidy in the diploid parent cannot easily be determined, because at least one diploid parent copy must be present to infer chromosomal inheritance patterns in triploid progeny. Further, these ratios are not fully informative because any copy number in the tetraploid parent (1/4, 2/4, 3/4, 4/4) can potentially exist in four observable cases in the triploid (4/3, 3/3, 2/3, 1/3). However, the percentage of reads attributable to the diploid parent in the triploid hybrid (i.e., percent diploid) can be utilized as a second parameter to rectify overlapping parent-hybrid ratios of different parent and hybrid combinations (Figure S1; Figure 3).

**Figure 3.**
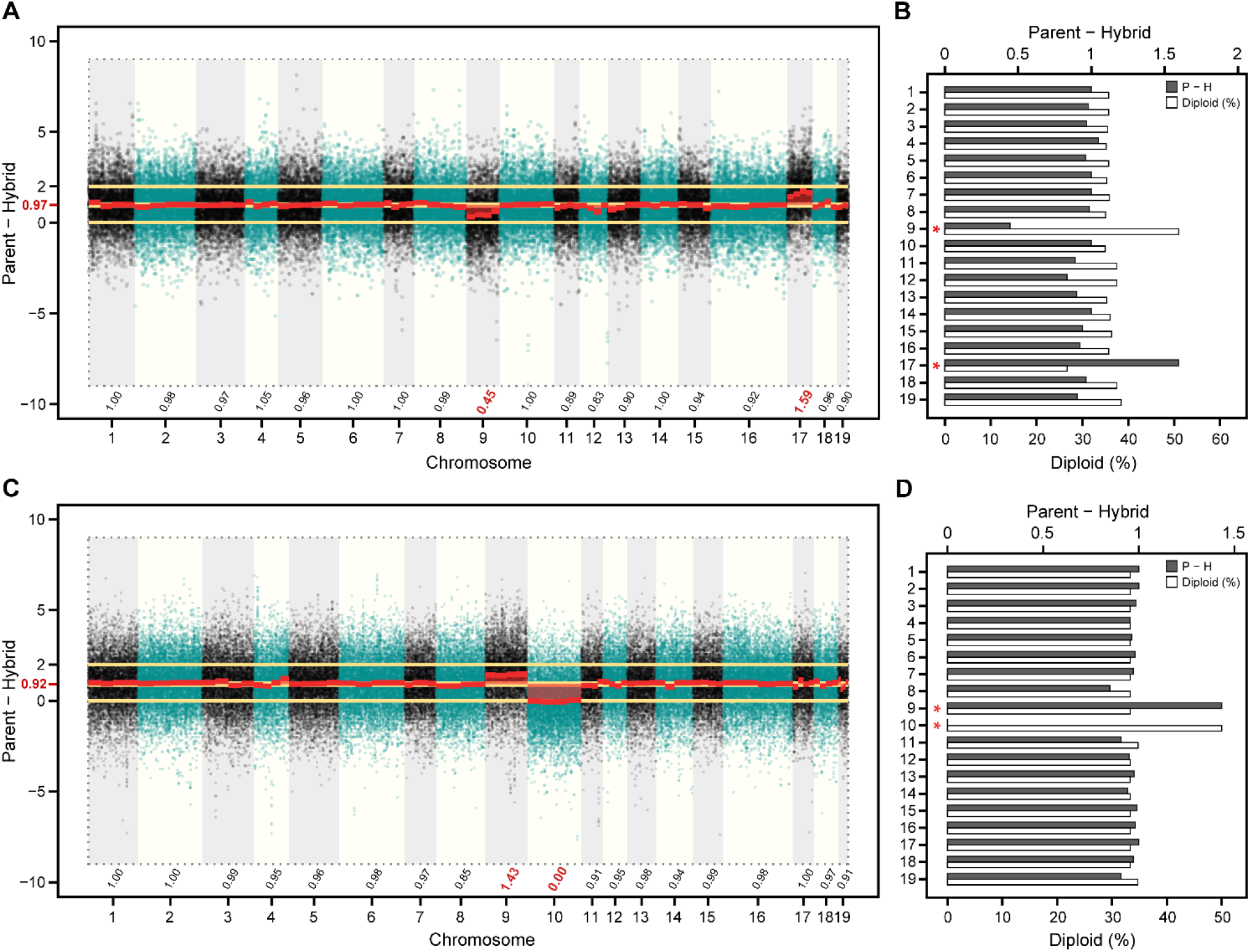
Manhattan plot **(A)** chromosome-wide differences of log_2_ (P_2X_ / P_4X_) expression (parent ‒ hybrid) between the family 415 parents (female diploid 94006 and male tetraploid 01-200-003) and the triploid hybrid 12X-415-031. Median parent ‒ hybrid values are shown above chromosome identifiers (x-axis). The barplot **(B)** depicts the median parent ‒ hybrid difference (dark grey bars, scale top x-axis) and the percent expression in the hybrid attributable to the diploid parent allele (white bars, scale lower x-axis) by chromosome (y-axis). The Manhattan plot in panel **(C)** and barplot in panel **(D)**, represent the same analyses, but between the family 423 parents (female diploid 07-MBG-5027 and male tetraploid 01-200-003) and the triploid hybrid 12X-423-070. Red text on x-axes in panels **(A)** and **(D)** correspond to red asterisks on y-axes in panels **(B)** and **(C)**, which denote significant differences (Wilcoxon *p*-value < 1×10^−16^).

While chromosome-wide log_2_ (P_2X_ / P_4X_) expression of the female diploid (*S. purpurea* 94006 and *S. viminalis* 07-MBG-5027) and male tetraploid (*S. miyabeana* 01-200-003) parents showed consistent median values approximately equal to 0, *Salix* chr09 significantly deviated from the expected (Wilcoxon *p*-value < 1×10^-16^), with a log_2_ (P_2X_ / P_4X_) of 0.49. This suggests that only three copies of chr09 were present in *S. miyabeana* parent 01-200-003. This was the case for both families 415 and 423, which share the male tetraploid parent 01-200-003. No significant deviations from the expected was observed for the parents of family 430 (*S. miyabeana* 01-200-006 × *S. viminalis* ‘Jorr’).

For family 415, five triploid individuals had a median log_2_ (P_2X_ / P_4X_) parent ‒ hybrid difference of 1.43 and approximately 34% of the reads which could be attributed to the diploid parent for chr09, which is expected, given the male parent was limited to three chr09 copies. The other five individuals had a log_2_ (P_2X_/P_4X_) difference of 0.45 for chr09 and were ∼50% diploid over all loci for the chromosome. Thus, the latter group in family 415 inherited two of the three tetraploid parent copies of chr09 and the former inherited only one. A total of six individuals in family 423 had a log_2_ (P_2X_ / P_4X_) difference of 1.44 and were 33.3% diploid on average for chr09, which is expected if they inherited two copies from the tetraploid, because family 415 and 423 share the same male tetraploid *S. miyabeana* parent. The other four individuals in family 423 had a log_2_ (P_2X_ / P_4X_) difference of 0.47 and were 50% diploid on average, so these individuals only inherited one of the three tetraploid parent copies of chr09.

It was not uncommon for individuals to possess an additional tetraploid copy of a chromosome and lack another. For instance, the family 415 individual, 12X-415-031, had a log_2_ (P_2X_ / P_4X_) difference of 1.59 for chr17, but only 25% diploid, which suggests that 12X-415-031 inherited an additional copy of the male tetraploid parent 01-200-003 chr17. Stunningly, the same individual also lacked one copy of the male chr09 (log_2_ (P_2X_ / P_4X_) = 0.45 and 50% diploid) (Figure 3A, Figure 3B). Another example was for the family 423 individual 12X-423-070 (Figure 3C, Figure 3D). While 12X-423-070 inherited two copies of chr09 from the tetraploid parent 01-200-003 (log_2_ (P_2X_ / P_4X_) = 1.43 and 33.3% diploid), this individual lacked one copy of the tetraploid parent chr10 (log_2_ (P_2X_ / P_4X_) = 0.0 and 50% diploid), which seems to be spurious, given there was no DNA-Seq or RNA-Seq coverage to indicate that 01-200-003 lacked a copy of chr10.

Unequal inheritance of chr09 in families 415 and 423 was unexpected, yet it permitted a test for genes insensitive to changes in dosage for this chromosome, as well as common genes up- or down-regulated in each group. Three individuals each from families 415 and 423 with a 33% diploid attribution and three each from both families with a 50% diploid attribution for chr09 were compared. Individuals with spurious tetraploid CN (e.g., 12X-415-031 and 12X-423-070) were not included in the analysis. As previously stated, there is a global dosage effect in triploids, irrespective of CN, but dosage sensitive genes, which are most likely to be misexpressed, should show consistent and directional fold-changes. To avoid any buffering effects from the diploid parent (P_2X_), allele-specific expression of P_4X_ in the parent and hybrid were compared with a binomial exact test to reject the null hypothesis that the expression of P_4X_ allele in the triploid hybrid is half (Pr = 0.5) that of P_4X_ allele in the tetraploid parent.

### Gene–trait correlations

Since CNV in triploids may posit drastic phenotypic consequences, Pearson correlations (*r*) were made for mean genome-wide diploid (%) and heterosis for important biomass-related growth traits collected in the field and greenhouse (Table S6). In general, diploid % was positively correlated with heterosis for foliar traits and inversely correlated with heterosis for biomass stem growth traits (Table 4; Figure 4) described in Carlson and Smart (2021). Diploid % was positively correlated with the field-collected leaf growth traits (length, perimeter, ratio, specific leaf area) and inversely correlated with stem growth traits (height, basal diameter, area, volume). For greenhouse-collected traits, diploid % was positively correlated with specific leaf area only, but inversely correlated with biomass yield, stem growth traits, and vegetative phenology. Field and greenhouse collected traits most positively correlated with diploid % were crown form (*r* = 0.65) and specific leaf area (*r* = 0.65), respectively, and inversely were plot height (*r* = −0.82) and root dry mass (*r* = −0.73), respectively. The only foliar field trait with an inverse relationship with diploid % was leaf shape factor (*r* = −0.51), which is a measure of leaf symmetry. Diploid % had a strong inverse relationship with the proportion of differentially expressed genes showing nonadditive inheritance (*r* = −0.71).

**Table 4.**
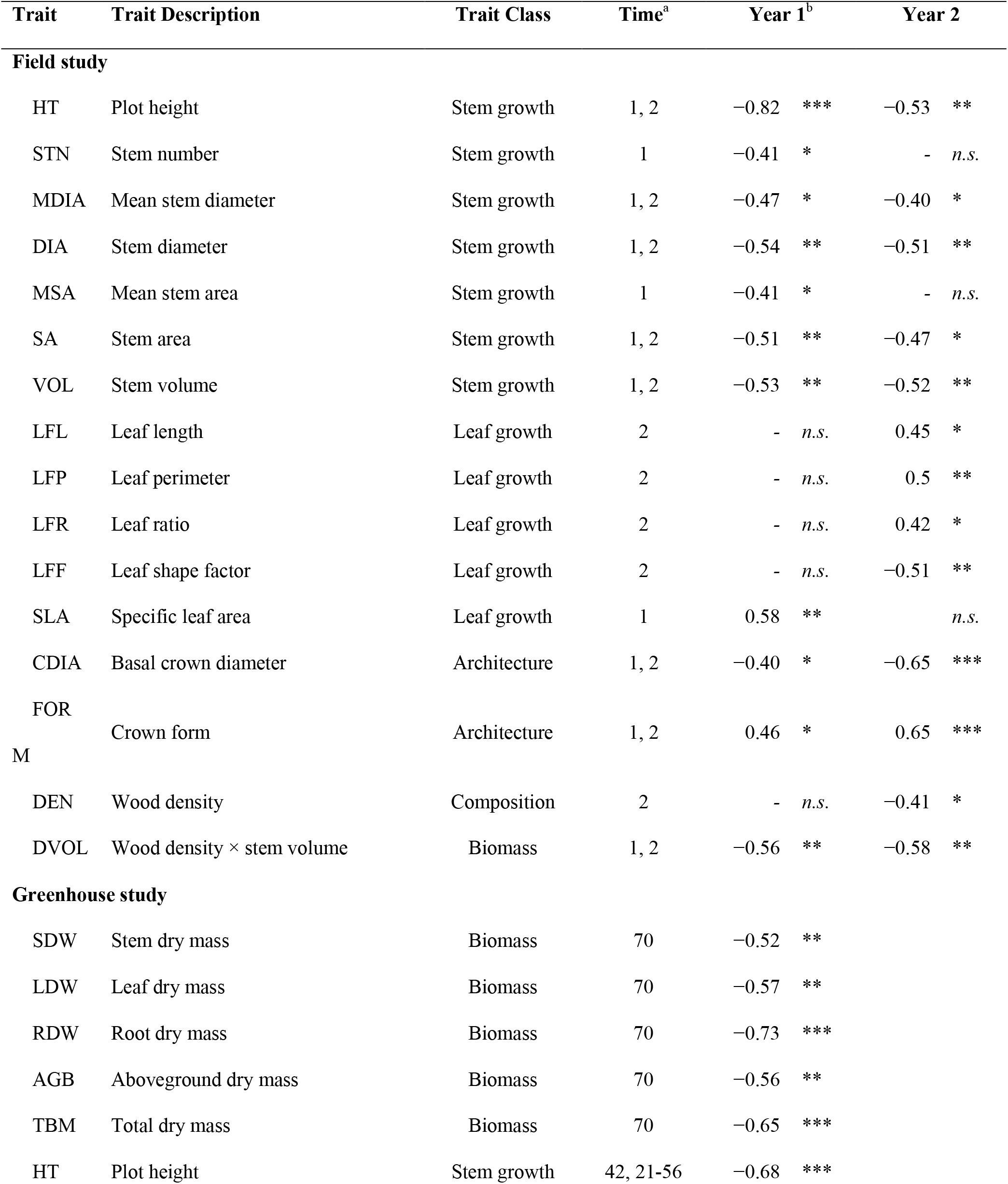
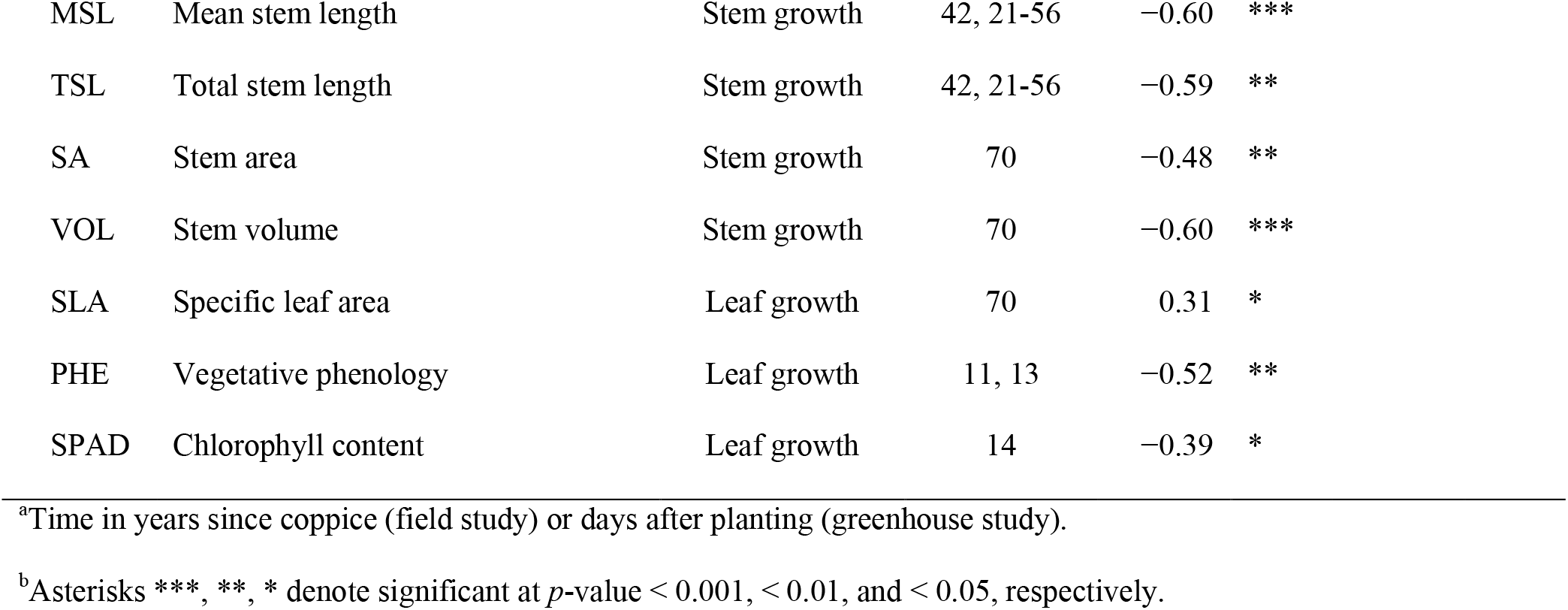
Pearson correlation coefficients (*r*) of heterosis for biomass-related traits and the mean percentage of each locus in triploid progeny attributable to the respective diploid parent (diploid %).

| Trait | Trait Description | Trait Class | Time<sup>a</sup> | Year 1<sup>b</sup> |  | Year 2 |  |
| --- | --- | --- | --- | --- | --- | --- | --- |
| Field study |  |  |  |  |  |  |  |
| HT | Plot height | Stem growth | 1, 2 | −0.82 | *** | −0.53 | ** |
| STN | Stem number | Stem growth | 1 | −0.41 | * | - | *n.s.* |
| MDIA | Mean stem diameter | Stem growth | 1, 2 | −0.47 | * | −0.40 | * |
| DIA | Stem diameter | Stem growth | 1, 2 | −0.54 | ** | −0.51 | ** |
| MSA | Mean stem area | Stem growth | 1 | −0.41 | * | - | *n.s.* |
| SA | Stem area | Stem growth | 1, 2 | −0.51 | ** | −0.47 | * |
| VOL | Stem volume | Stem growth | 1, 2 | −0.53 | ** | −0.52 | ** |
| LFL | Leaf length | Leaf growth | 2 | - | *n.s.* | 0.45 | * |
| LFP | Leaf perimeter | Leaf growth | 2 | - | *n.s.* | 0.5 | ** |
| LFR | Leaf ratio | Leaf growth | 2 | - | *n.s.* | 0.42 | * |
| LFF | Leaf shape factor | Leaf growth | 2 | - | *n.s.* | −0.51 | ** |
| SLA | Specific leaf area | Leaf growth | 1 | 0.58 | ** |  | *n.s.* |
| CDIA | Basal crown diameter | Architecture | 1, 2 | −0.40 | * | −0.65 | *** |
| FOR | Crown form | Architecture | 1, 2 | 0.46 | * | 0.65 | *** |
| M |  |  |  |  |  |  |  |
| DEN | Wood density | Composition | 2 | - | *n.s.* | −0.41 | * |
| DVOL | Wood density × stem volume | Biomass | 1, 2 | −0.56 | ** | −0.58 | ** |
| Greenhouse study |  |  |  |  |  |  |  |
| SDW | Stem dry mass | Biomass | 70 | −0.52 | ** |  |  |
| LDW | Leaf dry mass | Biomass | 70 | −0.57 | ** |  |  |
| RDW | Root dry mass | Biomass | 70 | −0.73 | *** |  |  |
| AGB | Aboveground dry mass | Biomass | 70 | −0.56 | ** |  |  |
| TBM | Total dry mass | Biomass | 70 | −0.65 | *** |  |  |
| HT | Plot height | Stem growth | 42, 21-56 | −0.68 | *** |  |  |

|  |  |  |  |  |  |
| --- | --- | --- | --- | --- | --- |
| MSL | Mean stem length | Stem growth | 42, 21-56 | −0.60 | *** |
| TSL | Total stem length | Stem growth | 42, 21-56 | −0.59 | ** |
| SA | Stem area | Stem growth | 70 | −0.48 | ** |
| VOL | Stem volume | Stem growth | 70 | −0.60 | *** |
| SLA | Specific leaf area | Leaf growth | 70 | 0.31 | * |
| PHE | Vegetative phenology | Leaf growth | 11, 13 | −0.52 | ** |
| SPAD | Chlorophyll content | Leaf growth | 14 | −0.39 | * |

<sup>a</sup>Time in years since coppice (field study) or days after planting (greenhouse study).
<sup>b</sup>Asterisks \*\*\*, \*\*, \* denote significant at $p$ -value $< 0.001$ , $< 0.01$ , and $< 0.05$ , respectively.

**Figure 4.**
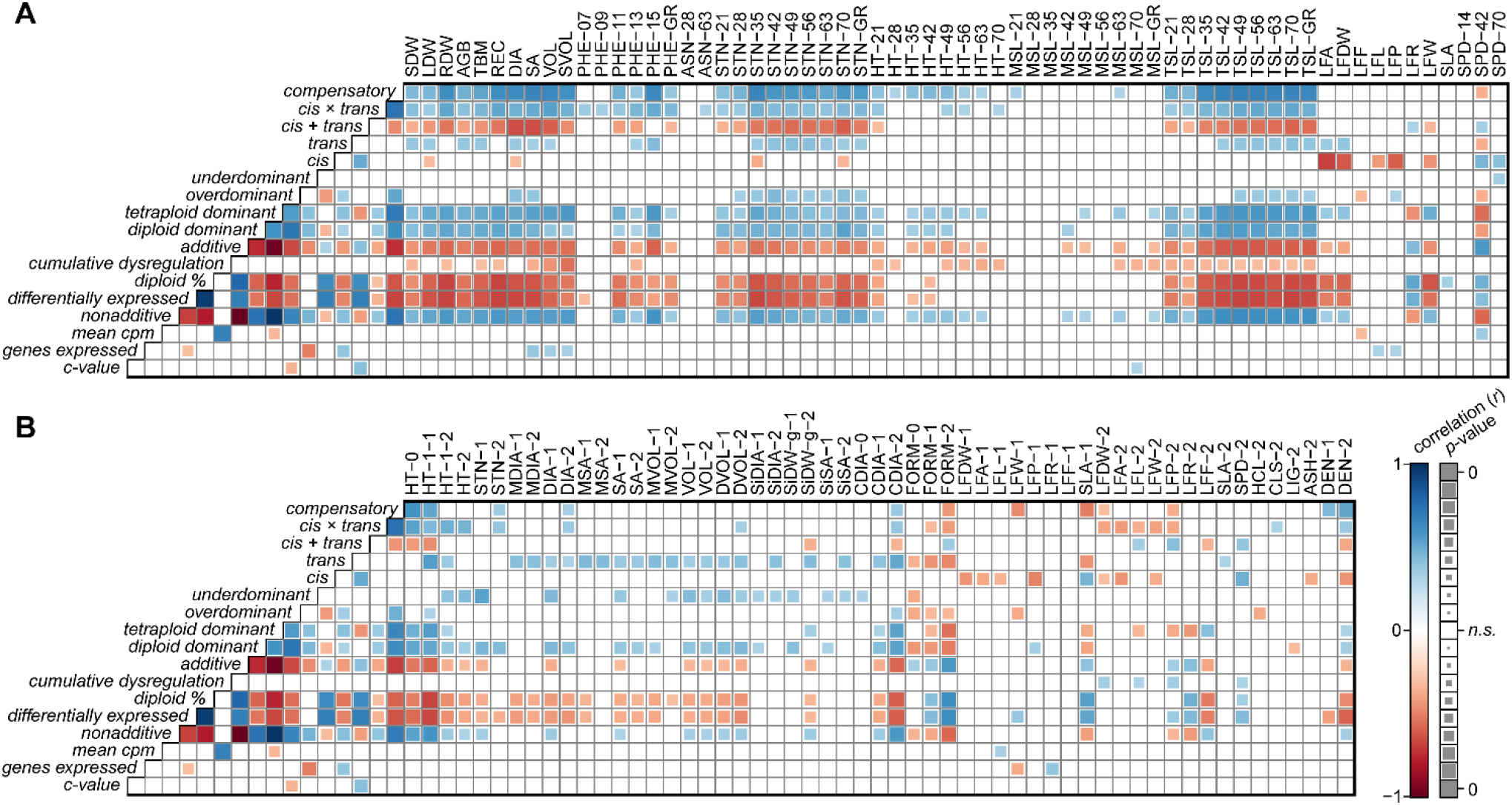
Correlations of nonadditive, regulatory divergent, and cumulative expression dysregulation with heterosis for **(A)** greenhouse and **(B)** field phenotypes. Pearson correlation coefficients (*r*), positive correlations are illustrated by filled blue squares and negative correlations by filled-red squares. Non-significant correlations (*p*-value > 0.01) were left blank. Significance levels (*p*-values) were used to scale the area of each square, such that smaller squares represent correlation coefficients with lower significance and larger squares represent those with greater significance.

Overall, there were stronger associations in the greenhouse trial than the field trial. The proportions of additive expression, and *cis*- and *cis* + *trans* divergence, were inversely correlated with heterosis for nearly all biomass traits in the greenhouse trial, as well as the total proportion of differentially expressed genes, cumulative expression dysregulation, and the proportion of differentially expressed genes with additive expression inheritance. For both the field and greenhouse trials, *trans*-divergence, differential expression, and the proportion of diploid- and tetraploid-parent dominant genes were positively correlated with heterosis for total stem volume. Heterosis for hemicellulose content was positively correlated with the proportion of *cis*-divergence and inversely with overdominance. Heterosis for cellulose and lignin content were positively and inversely correlated with the proportion of differentially expressed genes with diploid parent dominant expression. Chlorophyll content (SPAD) in both trials was inversely correlated with the proportion of nonadditive expression, *trans*, and compensatory regulatory divergence, and the proportion of differentially expressed genes with tetraploid parent dominant inheritance. *Cis*-divergence was positively correlated with the proportion of differentially expressed genes with additive inheritance, but inversely correlated with the proportion of diploid parent dominant and overdominant expression. The proportion of differentially expressed genes with *trans*-divergent expression was inversely correlated with the proportion of additive inheritance, but positively correlated with dominant and overdominant proportions. Finally, mean normalized expression levels (CPM) had the greatest positive association with cumulative dysregulation (*r* = 0.68); however, nonadditive inheritance and regulatory proportions lacked any significant associations with cumulative dysregulation.

Genes most commonly associated with heterosis for biomass growth traits were a peripheral-type benzodiazepine receptor (PBR, SapurV1A.0155s0220; *r* = 0.64 to 0.81) located on *Salix* chr02 and a squamosa promoter-binding-like protein (SPL10, SapurV1A.0056s0240; *r* = 0.67 to 0.73) on chr03 (File S3). Most common inverse gene associations with biomass growth traits were mediator of RNA polymerase II subunit 7 (MED7, SapurV1A.0616s0090; *r* = −0.74 to −0.81) on chr06 and a NLI interacting factor-like serine/threonine specific protein phosphatase (NIF, SapurV1A.0546s0050; *r* = −0.67 to −0.78) on chr15. Genes with a strong positive relationship with total biomass yield (*r* > 0.6, n = 189) were enriched for GO molecular functions: catalytic activity, transferase activity, and acyltransferase activity, and those inversely associated (*r* < −0.6, n = 94) were enriched for GO molecular functions: structural molecule activity and structural constituent of the ribosome (Table S7).

### Prediction of heterosis with random and selected gene expression sets

For most traits, genes that were differentially expressed between the F_1_ parents were more predictive of midparent heterosis than those 5,000 genes either randomly sampled or most highly expressed (Figure 5). Prediction accuracies of midparent heterosis using genes commonly differentially expressed between all three family parent pairs (n = 379) were akin to the larger set of genes differentially expressed between at least one pair (n = 4,986), yet there were cases in which one gene set performed substantially better, vice versa.

**Figure 5.**
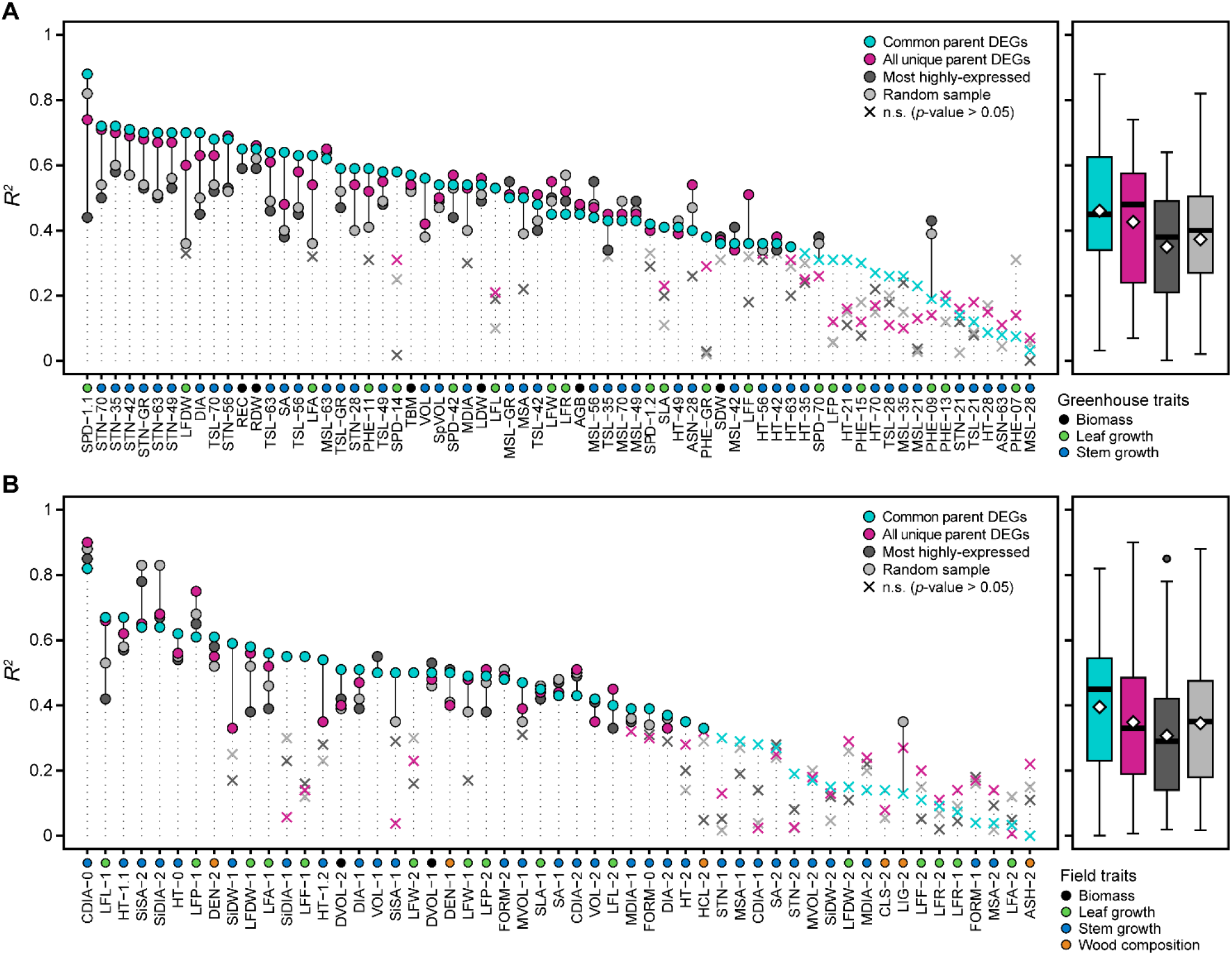
Prediction accuracies (*R*^2^) of heterosis values for **(A)** greenhouse and **(B)** field phenotypes using selected and random gene expression sets. These four gene sets were: differentially expressed genes common among parent pairs (cyan, n = 379), differentially expressed genes in at least one parent pair (magenta, n = 4,978), the most highly expressed genes (dark grey, n = 5,000), and a random sampling of genes (light grey, n = 5,000). Boxplots to the right of each panel depict the distribution of prediction accuracies, with means represented as yellow diamond points. Traits are colored according to respective classes described in the lower left legend of each panel.

Overall, mean prediction accuracies were greater for greenhouse traits (Figure 5A) than field traits (Figure 5B), of which the common, overlapping differential expression gene set had better prediction accuracies on average. This can partly be explained by more frequent repeated measurements recorded in the greenhouse and the impact of pest damage on field phenotypes not present in the greenhouse. In addition, lower *R*^2^ values were observed for second year post-coppice measurements in the field compared to those taken the first year following coppice. Across all four gene sets, basal crown diameter had the highest prediction accuracies (*R*^2^ = 0.82 – 0.90), followed by leaf perimeter, chlorophyll content, primary stem number, and basal stem diameter.

## DISCUSSION

### Differential gene expression is both additive and nonadditive

Using microarrays of maize, Stupar and Springer (2006) determined that approximately 20% of the genes that were differentially expressed between inbred parents were nonadditively expressed in the hybrid, although very few were above the high parent (overdominant) or below the low parent (underdominant). Swanson and Wagner (2006) found all inheritance categories in the hybrid represented among differentially expressed genes between two inbred parents of maize. In our study, there were very few genes that were differentially expressed between diploid and tetraploid parents and were outside the parental range in the triploid hybrid, especially overdominant genes, which were three-times less-frequent on average than underdominant genes. In contrast, differentially expressed genes between heterozygous thistle (*C. arvense*) parents were more frequently overdominant than underdominant in intraspecific hybrids (Bell *et al*. 2013). Expression-level dominance was most-prominent in both shoot tip and stem internode tissues of F_1_ and F_2_ diploid *S. purpurea* families (Carlson *et al*. 2017) and was primarily biased in the direction of the female parent, especially in shoot-tip tissues. Very little additive gene expression was observed in *S. purpurea*, which is a unique result, compared with model crop plants (Guo *et al*. 2006; Stupar and Springer 2006; Song *et al*. 2013).

Among the triploid *Salix* families investigated in this study, expression-level dominance was prominent, as was established in diploid *S. purpurea*, but the percentage of differential expression attributed to dominance inheritance ranged from 28% to 60%. Progeny from reciprocal crosses between *Salix* Sections Vimen and Helix, showed the greatest percentage of dominant expression, which was 50% of those genes expressed differently between diploid and tetraploid parents. Cases of expression-level dominance in polyploid crops have been described in intraspecific thistle (Bell *et al*. 2013), interspecific coffee (Combes *et al*. 2015), as well as in allotetraploids of both rice (Xu *et al*. 2014) and Arabidopsis (Shi *et al*. 2012). This preferential expression is thought to be orchestrated by allelic interactions, which functions to silence one of the parent alleles in a parent-of-origin manner (Chen and Pikaard, 1997; Stupar *et al*. 2007; Donoghue *et al*. 2014; Baldauf *et al*. 2016).

Further analysis of allele-specific expression in triploid hybrids of willow indicated that gene expression variation was associated with both *cis*- and *trans*-regulatory divergence, and that *cis*‒*trans* compensatory interactions accounted for up to 25% of the variation. Allele-specific expression has been extensively studied in model species, most notably, in interspecific hybrids and allopolyploids of Arabidopsis (Shi *et al*. 2012) and *Drosophila* (Landry *et al*. 2005; Wittkopp *et al*. 2008a; Wittkopp *et al*. 2008b; McManus *et al*. 2010). There is a general trend that *cis*-regulatory divergence accounts for a greater proportion of expression variation in interspecific hybrids and that *trans*-regulatory divergence is more frequent in intraspecific hybrids (Wittkopp *et al*. 2004). In hybrids of inbred maize, *cis*-acting variation accounted for most of the divergent expression between parents and was largely attributed to additive expression patterns (Stupar and Springer 2006). Greater sequence divergence was proposed to promote the flexibility of *trans*-factors in their binding to interacting factors and *cis*-elements in *Arabidopsis thaliana* and *A. arenosa* parent alleles (Shi *et al*. 2012). McManus et al. (2010) hypothesized that greater transgressive inheritance is associated with greater proportions of *cis* × *trans* divergence. Likewise, what was identified in triploid *Salix* hybrids, greater proportions of overdominant and underdominant (transgressive) expression inheritance did coincide with greater proportions of *cis* × *trans* and compensatory regulatory classes. Further, a greater proportion of *cis* + *trans* divergence coincided with a lower proportion of underdominant expression.

### Global dosage balance with local sensitivities

Dosage in all three triploid families appeared to behave in an extraordinarily additive manner, irrespective the number of parent copies inherited. However, a handful of genes did depart from expected dosage in triploids, most notably, those coding for heat shock proteins. In this study, genes annotated as coding for heat shock proteins displayed greater expression in individuals with normal chr09 copies, whereas those null for a tetraploid parent copy had greater expression of stress- or senescence-associated genes. Overall, there were greater proportions of loci showing *cis* × *trans* and compensatory regulatory patterns in family 415 and 423 individuals that were aneuploid with only one tetraploid parent copy (e.g., chromosomes 2, 9, 10, and 17), or a greater average diploid %. While the quantity of a translation product (protein subunit) may impact the assembly of a particular complex, the mere involvement in a complex can also impact protein stability (Veitia *et al*. 2007). It may be that null mutations in metabolic functions are tolerated in a heterozygous state, but only weak, loss-of-function, dosage-sensitive genes can survive negative selection as heterozygotes (Birchler and Veitia 2010). The balance of regulatory hierarchies (dosage balance) (Birchler *et al*. 2005) are sensitive to gene dosage and changes in individual components can influence phenotype. In macromolecular complexes, dosage balance is essential, because partial aneuploidy of a dosage-sensitive gene can change the stoichiometry of the complexes and lead to fitness defects (Veitia *et al*. 2008). In maize, greater proportions of nonadditive expression was observed in triploid and tetraploid hybrids with genome dosage effects (Guo *et al*. 1996; Auger *et al*. 2005; Birchler *et al*. 2005; Riddle *et al*. 2010).

Previous gene expression studies in inbred and outcrossing species have regularly pooled F_1_ progeny libraries prior to sequencing. While this is not an issue for inbred crops, out results show that pooled RNA-Seq can underestimate factors contributing to the inheritance of gene expression in heterozygous species, especially for families derived from natural polyploids. For instance, without sequencing individual libraries, we would not have detected aneuploidy for chromosomes of polyploid progenitors in the F_1_ based on pooled RNA-Seq data alone, which could distort assumptions about the evolution of gene expression inferred from inheritance and regulatory assignments. Even if the expected ploidy in the hybrid is based on chromosome counts or DNA-Seq of the parents, there may not be equal inheritance, and binomial tests for ASE between the parents and the hybrid would be incorrect if based on a fixed probability estimate. Thus, prior to tests for ASE, a simple adjustment could be made, which would first require that each chromosome (or scaffold) be tested independently. Utilizing median fold-changes in the parents and the percentage of reads in the hybrid attributable to the diploid parent, a probability of success under the null could be properly assigned.

Beyond the fact that the parents in this study were highly heterozygous, it is possible that CNV or aneuploidy can help explain some of the variation in heterosis observed within and among triploid families. In aneuploid studies, changing numerous chromosome segments can alter quantitative characters (Guo and Birchler 1994). Here, genome-wide averages of ASE attributable to the diploid parent (diploid %) in triploids was inversely correlated with heterosis for important stem growth traits (e.g., total harvestable biomass), but positively with foliar traits. The dosage balance hypothesis, outlined by Birchler (2005), may very well apply to slight deviations in the global inheritance of parent ASE or major differences in chromosomal copy number, as was observed for *Salix* chr09 aberrations in families 415 and 423. Genetic mapping in F_2_ *S. purpurea* (Carlson *et al*. 2019) identified QTL on chr09 for leaf length, leaf perimeter, and specific leaf area, so positive correlations between diploid % and foliar traits could indicate a dosage sensitivity of genes controlling the variation for these traits.

A cluster of genes collocated on a 50 kb interval on chr10 contained genes highly-expressed in tetraploid *S. miyabeana* parents and triploid progeny, but with very low expression in diploid parents. These genes included duplicates, annotated as 3’-N-debenzoyl-2’-deoxytaxol N-benzoyltransferase (DBTNBT, TAX10), which catalyzes the final step in biosynthesis of the anti-cancer compound Taxol (Walker *et al*. 2002). The constitutive high expression levels of DBTNBT in triploids and tetraploid *S. miyabeana* parents could suggest involvement in the synthesis of an important defensive compound in *Salix*. There are multiple copies of genes with this annotation in the *Salix* genome and a number of them collocate with QTL for variation in tremuloidin on chr08, 10, 15, and 16 (Keefover-Ring *et al*., in preparation). The abundance of tremuloidin or a related phenolic glycoside could be a source of broad-spectrum pest and/or disease resistance conferred to triploid hybrids by *S. miyabeana* parents, as all triploid genotypes in Carlson and Smart (2021) displayed field resistance to most willow pests and pathogens, but not intra- and interspecific diploids. In a F_2_ *S. purpurea* mapping population, Carlson *et al*. (2019) identified QTL associated with many important traits for biomass production. One QTL for willow leaf rust (*Melampsora* spp.) incidence was identified on chr10, and DBTNBT genes described here fall within that confidence interval, meriting further investigation.

### Nonadditive gene expression correlates with nonadditive phenotypic expression

One of the major challenges in molecular genetics is disentangling the relationship of transcriptome-wide expression patterns to phenotypic effects (Birchler *et al*. 2007). Rather than concentrating on the terminologies of heterosis models (e.g., dominance, overdominance, or pseudo-overdominance), Birchler (2010) promoted a progression to a more nuanced quantitative and interactive or network-oriented framework for dissecting the phenomenon of heterosis. We utilized DNA-Seq and RNA-Seq to unravel the underlying regulatory architecture of differential expression and improve our understanding of heterosis in high-yielding triploid hybrids of willow. We showed that the proportion of genes differentially expressed between diploid and tetraploid parents attributable to nonadditive gene expression in the triploid hybrid (namely expression-level dominance) was positively correlated with heterosis for biomass yield as well as biomass-related growth traits collected in the greenhouse and in the field. In addition, we corroborate some of the key findings reported in Kremling *et al*. (2018), for example that cumulative expression dysregulation is inversely correlated with heterosis for biomass; and that individuals with greater absolute expression tended to display greater levels of dysregulation.

Importantly, tetraploid parent dominant genes among triploid hybrids were enriched for the following pathways: phenylpropanoid biosynthesis, cyanoamino acid metabolism, biosynthesis of secondary metabolites, and starch and sucrose metabolism. Some of the most intriguing tetraploid parent dominant genes identified in this study were those annotated as uridine diphosphate (UDP) glycosyltransferases (UGTs). UGTs catalyze the transfer of sugars to a wide range of acceptor molecules, including plant hormones, and all classes of plant secondary metabolites (Ross *et al*. 2001). Both tetraploid parent dominant genes and those genes positively correlated with biomass yield and stem growth traits were enriched for the GO molecular function of catalytic activity. Further analysis of these candidate genes and gene sets, with regards to their relevance in overlapping support intervals from mapping experiments or regulatory patterns in other high-yielding triploid hybrid individuals, will prove useful in the genetic improvement of shrub willow as a bioenergy crop.

### Parent differentially expressed genes are most predictive of heterosis in F_1_ hybrids

Here, we tested whether parent differentially expressed genes are predictive of heterosis in three interspecific triploid F_1_ *Salix* crosses, by comparing prediction accuracies of those gene sets to a random sampling of genes as well as a selection of genes most highly expressed. While it would be assumed that the most highly expressed genes would also be the most variable, these genes had the lowest mean prediction accuracies for most traits and performed similarly as did a random sampling of genes of equal size. Differentially expressed genes were most predictive of heterosis, and often moreso using a reduced gene set of only common, overlapping differentially expressed genes among family parent pairs. However, there were a handful of traits where prediction accuracies of all gene sets were not considerably different from that of a random sample. This could mean that midparent heterosis values are attributable to population structure and/or highly quantitative in nature, such that a random gene sample is sufficient to illustrate the inherent transcriptome-wide differences between parent species. Thus, strong family-specific responses to hybridization and transgressive phenotypic expression would result in higher prediction accuracies for specific traits that have high among but not within family variances. Yet, this was not often the case. The phenotypes used in this study were from Carlson and Smart (2021), which reported both hybrid vigor and hybrid necrosis within all intra- and interspecific F_1_ families in field and greenhouse conditions. Midparent heterosis values were normally distributed, besides traits with low variance, like later vegetative phenology dates.

Genes that were differentially expressed between the parents showed primarily additive and dominant inheritance patterns among F_1_ progeny, but segregated within families. Progeny individuals with a greater frequency of genes with dominant expression patterns were more apt to display heterosis for biomass growth. Further, genes that were differentially expressed between parents were more predictive of heterosis in F_1_ progeny compared to a random sampling of genes, irrespective of expression level. These gene sets could be used to aid in the selection of genotypes or breeding populations in the greenhouse by utilizing expression levels as an indicator of performance based on prior related datasets. While only three species were assayed in this study, the inclusion of additional, diverse parent species of varying heterozygosity would help determine if there is a core set of genes and/or transcriptional regulators that, when differentially expressed, comprise a network predictive of triploid heterosis in F_1_ crosses.

This work highlights regulatory factors influencing differential expression, as well as genes and gene sets predictive of heterosis for biomass growth, physiological, and wood chemical composition traits collected in the greenhouse and field. It is vital that we apply our ever-improving understanding of heterosis from studies of well-characterized diploid crop species, such as maize, tomato, and rice to the improvement of yield and biomass quality of undomesticated crops, including willow and poplar, which provide sustainable sources of lignocellulosic biomass for bioenergy, biofuels, and bioproducts. Additional characterization of the genomic basis of heterosis in related genera or more diverse *Salix* crosses will be valuable in understanding the broad evolutionary benefits of wide hybridization and incidence of polyploidy.

## DATA AVAILABILITY STATEMENT

The gene expression data (File S1), heterosis values (File S2), and gene-trait correlations (File S3) used in this paper, as well as Supplementary Tables S1-S7 and Figures S1-S4 are available online: www.github.com/Willowpedia/Carlson2021_TriploidHeterosis.

## ACKNOWLEDGEMETS

We are grateful to Lauren Carlson and Dawn Fishback for their excellent technical assistance, and to Dr. Fred Gouker and Dr. Eric Fabio for their assistance with harvest, processing, and tissue collection.

## AUTHOR CONTRIBUTIONS

CHC wrote the manuscript, performed DNA and RNA isolations, phenotyping, bioinformatics, and statistics, YC and APC conducted sequencing and bioinformatics, CT and LBS devised the study and managed research programs. All authors participated in reviewing and editing the manuscript.

